# Building a robust chromatin immunoprecipitation (ChIP) method with substantially improved efficiency

**DOI:** 10.1101/2020.02.20.958330

**Authors:** Huimin Zhao, Hongyan Li, Yaqi Jia, Xuejing Wen, Huiyan Guo, Hongyun Xu, Yucheng Wang

## Abstract

Chromatin immunoprecipitation (ChIP) is the gold-standard method to detect the interactions between proteins and chromatin, and is a powerful tool to identify epigenetic modifications. Although ChIP protocols for plant species have been developed, many specific features of plants, especially woody plants, still hinder the efficiency of immunoprecipitation, resulting inefficient ChIP enrichment. There is an active demand for a highly efficient ChIP protocol. In the present study, we employed *Betula platyphylla* (birch) and *Arabidopsis thaliana* as the research materials, and five factors closely associated with ChIP efficiency were identified, including crosslinking, chromatin concentration using centrifugal filter, using new immunoprecipitation buffer, rescue DNA with proteinase K, and using sucrose to increase immunoprecipitation efficiency. Optimization of any these factors can significantly improve ChIP efficiency. Considering these factors together, a robust ChIP protocol was developed, for which the average fold enrichments were 16.88 and 6.43 fold of that gained using standard ChIP in birch and Arabidopsis, respectively. As this built ChIP method works well in both birch and Arabidopsis, it should be also suitable for other woody and herbaceous species. In addition, this ChIP method make it is possible to detect low-abundance TF-DNA interactions, and may extend the application of ChIP in plant kingdom.

**One sentence summary:** Building a ChIP method that increases fold enrichment of birch by 16 folds in average and is adapted for both woody and herbaceous plants.

## INTRODUCTION

Chromatin immunoprecipitation (ChIP) is an important technique that is widely used to examine epigenetic modifications or identify protein-DNA interactions. Transcription factors (TFs) bind to regulatory sequences to modulate gene expression, and gene expression regulation plays an essential role in various of cellular processes. However, which genes are directly regulated by the TFs, and how the TFs control gene expression *in vivo*, remain largely unknown. ChIP is a powerful tool to identify genes that are regulated directly by certain TFs and to address TFs’ recognition of their target genes *in vivo*. In addition, combined with high throughput sequencing technology, ChIP-seq has become the gold-standard method to detect binding regions for TFs on a genome-wide scale (Verkest *et al.*, 2014). However, current ChIP procedures have shortcomings in terms of the overall inefficiency of ChIP enrichment, making it difficult to detect low-abundance TF-DNA interactions (Verkest *et al.*, 2014).

Although ChIP protocols for plant species have been developed (Bowler *et al.*, 2004, Gendrel *et al.*, 2005, Saleh *et al.*, 2008, Kaufmann *et al.*, 2010), some specific features of plants species, such as rigid cell walls, chloroplasts, the paucity of nuclei in some tissues, and large vacuoles, all markedly hinder the immunoprecipitation of DNA and represent a challenge for TF-DNA enrichment. Therefore, genome-wide ChIP studies of plant species are lagging behind those of other eukaryotic systems (Verkest *et al.*, 2014). In addition, woody plants have thick-walled cells, and high levels of phenolics and/or polysaccharides, which adversely affect many key steps in ChIP procedures that have been optimized for tissues or cells of non-woody plants (Li *et al.*, 2014). An alternative approach to ChIP, Chromatin Affinity Purification (ChAP), had been proposed, which does not require immunoprecipitation, and is effective in plant chromatin studies (Zentner et al., 2014). Additionally, tandem chromatin affinity purification (TChAP) has also been developed in *Arabidopsis thaliana* plants, which can greatly improve DNA enrichment efficiency compared with ChIP (Verkest *et al.*, 2014). However, both ChAP and TChAP cannot be used in epigenetic studies, such as post-translational histone modifications, and standard ChIP is the most used method in epigenetics. At the same time, in the study of TF-DNA interactions, standard ChIP is still widely used in most case studies because of its properties. Therefore, standard ChIP is an important tool in molecular biology investigations. Standard ChIP does not work well in some plant species, especially woody plants; therefore, there is an active demand to overcome the inefficiency of ChIP enrichment, and to improve standard ChIP to make it suitable for both woody and herbaceous plants.

In the present study, we studied the factors that influence the efficiency of standard ChIP, and some key processes in the ChIP protocol were identified and optimized. Consideration of the factors involved in ChIP efficiency allowed us to develop a ChIP method that could significantly improve ChIP efficiency and could work well in woody and herbaceous plants. In addition, the results obtained in the present study could be used to develop an efficient ChIP protocol for use in other eukaryotic species, and the strategies and technologies used to optimize ChIP could also be used in other techniques that involve immunoprecipitation.

## RESULTS

### Crosslinking using 3% formaldehyde can resist decrosslinking caused by sonication better than 1% formaldehyde

Decrosslinking of chromatin during ChIP will reduce the yield. Sonication treatment can cause decrosslinking of chromatin; therefore, we first studied whether different concentrations of formaldehyde treatment could affect the decrosslinking caused by sonication. Birch plants (*Betula platyphylla*) were crosslinked using 1% and 3% formaldehyde, respectively, and then treated with sonication for chromatin fragment. After sonication, the decrosslinked DNA was harvested by extraction with Tris-phenol and chloroform, and analyzed using quantitative real-time polymerase chain reaction (qPCR). The results showed that 3% formaldehyde treatment could reduce the decrosslinking of chromatin by 0.7–2.98 fold compared with 1% formaldehyde treatment after sonication (Fig. 1). Therefore, considering the effects of decrosslinking chromatin caused by sonication, it is better to crosslink chromatin and proteins using 3% formaldehyde rather than 1% formaldehyde.

**Figure 1.**
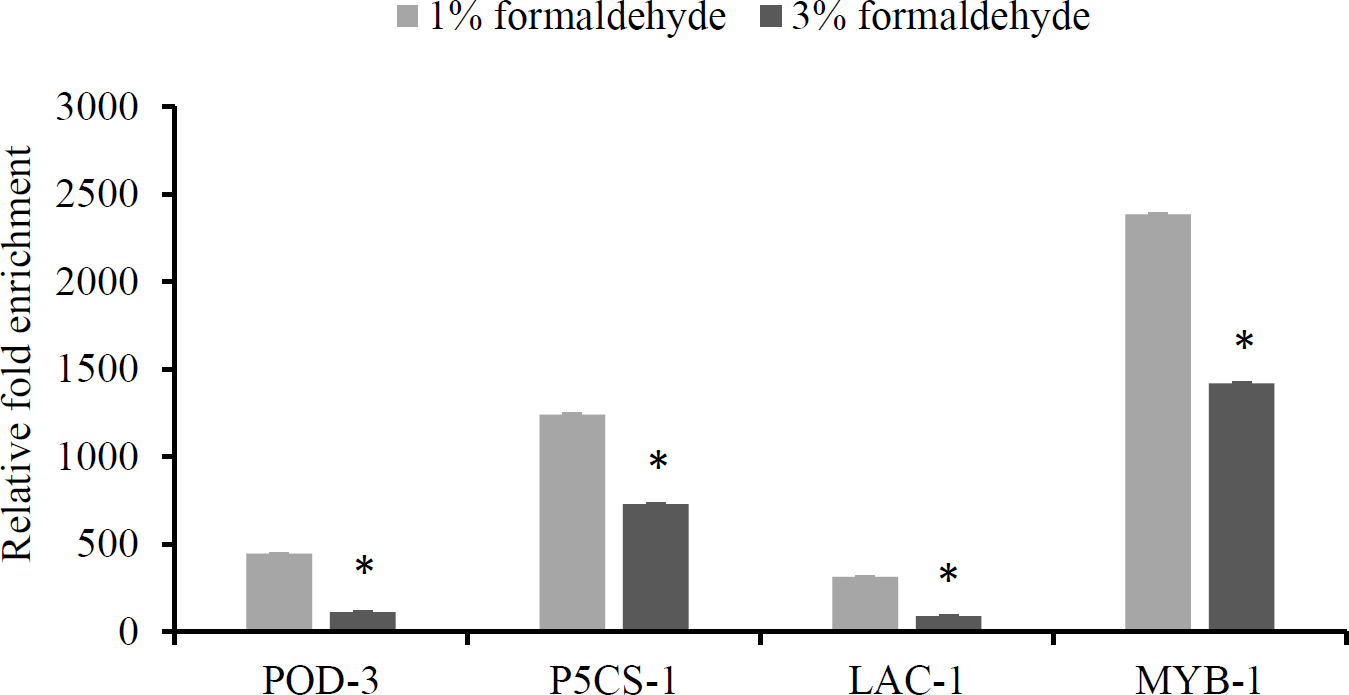
Comparison of decrosslinking caused by sonication between chromatin crosslinked by 1% and 3% formaldehyde. Four truncated promoters from birch that had been confirmed to be directly bound by BplMYB46 were studied. The chromatin was crosslinked with 1% or 3% formaldehyde, and treated with sonication. After sonication, the de-crosslinked DNA was harvested by extraction with Tris-phenol and chloroform, and analyzed using qPCR. Three independent experiments were performed, and data are means ± SD from three replicates.

### The concentration of the crosslinked chromatin significantly increases the immunoprecipitation efficiency

To study the effects of the concentration of formaldehyde-crosslinked-chromatin on immunoprecipitation, the sonicated chromatin from transgenic birch overexpressing *BplMYB46* was purified and concentrated using a 30 kDa cutoff centrifugal filter, and then the same quantity of antibody for immunoprecipitation as standard ChIP method was used. The standard ChIP procedure (described in the section of “Procedure for standard ChIP”) was also performed as the control. ChIP-qPCR was performed to check the efficiency of the enrichment of ChIP products. Four genes whose promoters had been confirmed to be bound by BplMYB46 in birch were analyzed. After the chromatin was concentrated using protein centrifugal filters, the enrichment increased markedly to 2.01 to 3.31-fold higher than that gained without using the centrifugal filter (Fig. 2). These results suggested that concentration of the crosslinked chromatin using a protein centrifugal filter is quite important for immunoprecipitation, resulting in a significant increase in immunoprecipitation efficiency and improving the enrichment of ChIP DNA.

**Figure 2.**
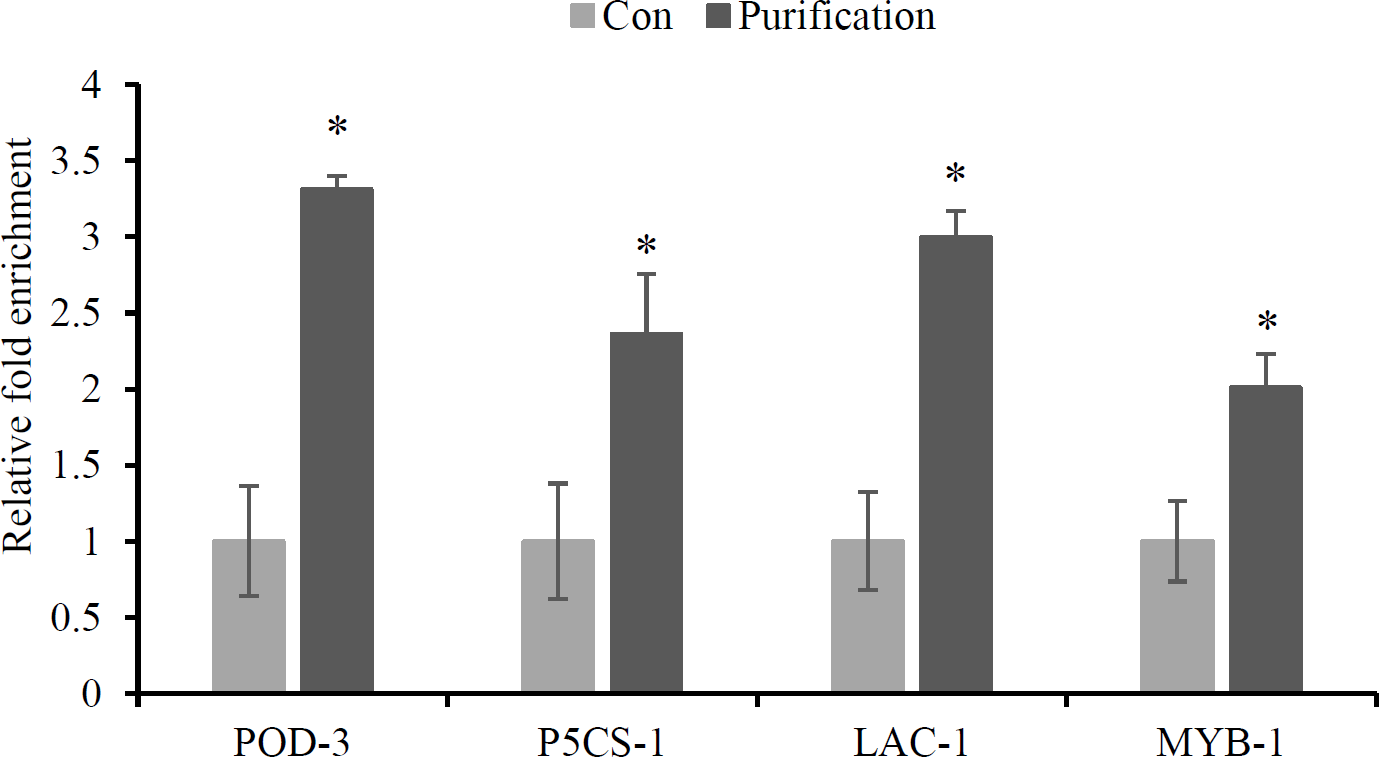
Concentration of chromatin using protein centrifugal filters significantly improves ChIP efficiency. Formaldehyde crosslinked chromatin was purified and concentrated using protein ultrafiltration centrifugal tube, and then was used for immunoprecipitation. Con: Fold enrichment in the standard ChIP protocol. Concentration: Fold enrichment of ChIP performed with concentrated chromatin with centrifugal filter. The relative fold enrichment was calculated as: Concentration/Con. Four truncated promoters of genes from *B. platyphylla* that were previously confirmed to be directly regulated by *BplMYB46* were analyzed, and their fold enrichments were determined using ChIP-qPCR. Three independent experiments were performed, and data are means ± SD from three replicates.

### A new buffer for immunoprecipitation in ChIP

The immunoprecipitation buffer is very important for the immunoprecipitation and enrichment of ChIP DNA. To optimize the immunoprecipitation buffer, we developed a new buffer, termed optimized immunoprecipitation (OIP) buffer, which has a similar pH value (pH = 7.4) to that of the plant nucleus (Shen et al., 2013). ChIP was performed using the OIP buffer, and the classic IP buffer (ChIP Ab incubation buffer) was used as a control. The ChIP-qPCR results showed that using the OIP buffer improved the efficiency of ChIP significantly, with an enrichment of 2.58–4.72-fold compared with that gained using the ChIP Ab incubation buffer (Fig. 3). These results suggested that the OIP buffer is more efficient for immunoprecipitation than the ChIP Ab incubation buffer.

**Figure 3.**
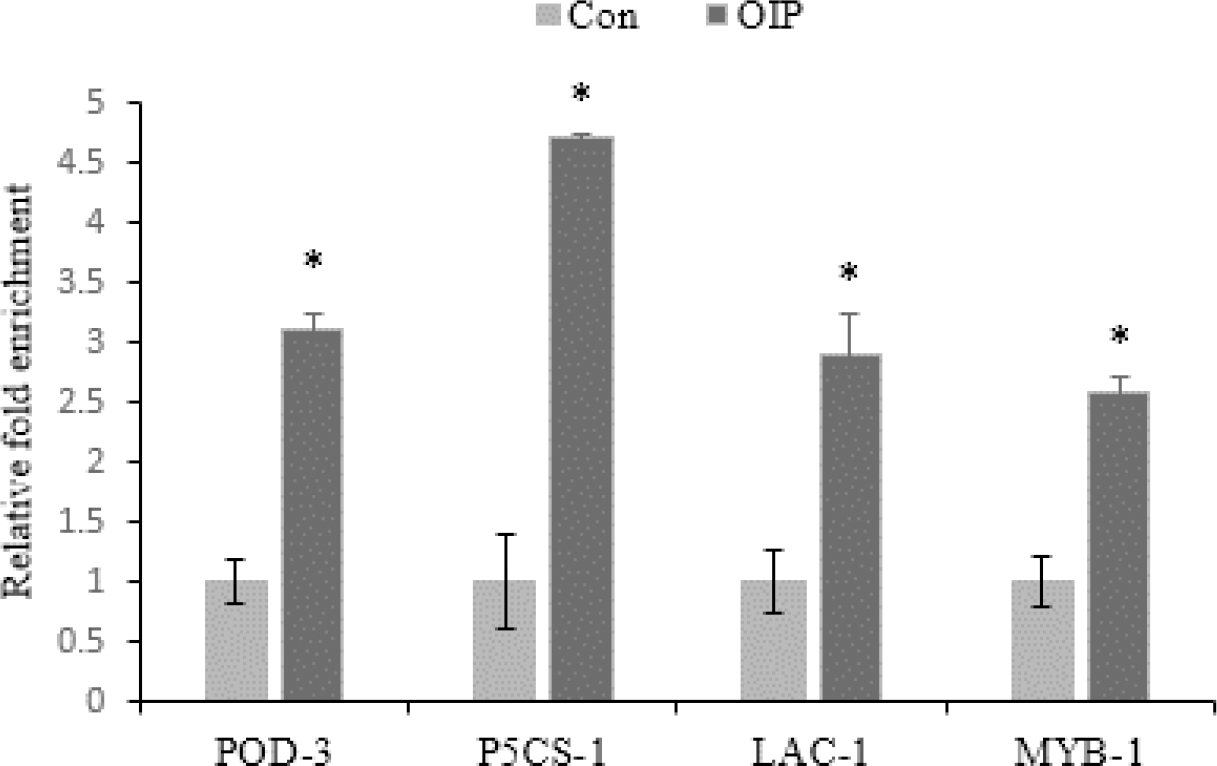
Determination of ChIP efficiency using the optimized immunoprecipitation buffer. The OIP (optimized immunoprecipitation) buffer was used for immunoprecipitation, and the immunoprecipitation buffer from the standard ChIP protocol was used as a control. Con: Fold enrichment of ChIP performed using ChIP Ab incubation buffer from the classic protocol, which was used as control; OIP: Fold enrichment of ChIP performed using the OIP buffer. The relative fold enrichment was calculated as: OIP/Con. Four truncated promoters of genes directly regulated by *BplMYB46* were studied in *B. platyphylla* using ChIP-qPCR. Three independent experiments were performed, and data are means ± SD from three replicates.

### Determination of the most suitable NaCl concentration for immunoprecipitation

Previous research showed that 150 mM NaCl is not a suitable concentration for immunoprecipitation (Li *et al.*, 2014); therefore, we determined whether 150 mM NaCl is a suitable concentration for immunoprecipitation in the OIP buffer. OIP buffers with NaCl at 100, 120, 150, and 170 mM were used, and the concentrations of the other reagents in the OIP buffer are unchanged. The results showed that 150 mM NaCl displayed highest immunoprecipitation efficiency, followed by 170 mM NaCl; however, 100 and 120 mM NaCl showed relatively low immunoprecipitation efficiency (Fig. 4). This result suggested that 150 mM NaCl is the most suitable concentration for immunoprecipitation in the OIP buffer.

**Figure 4.**
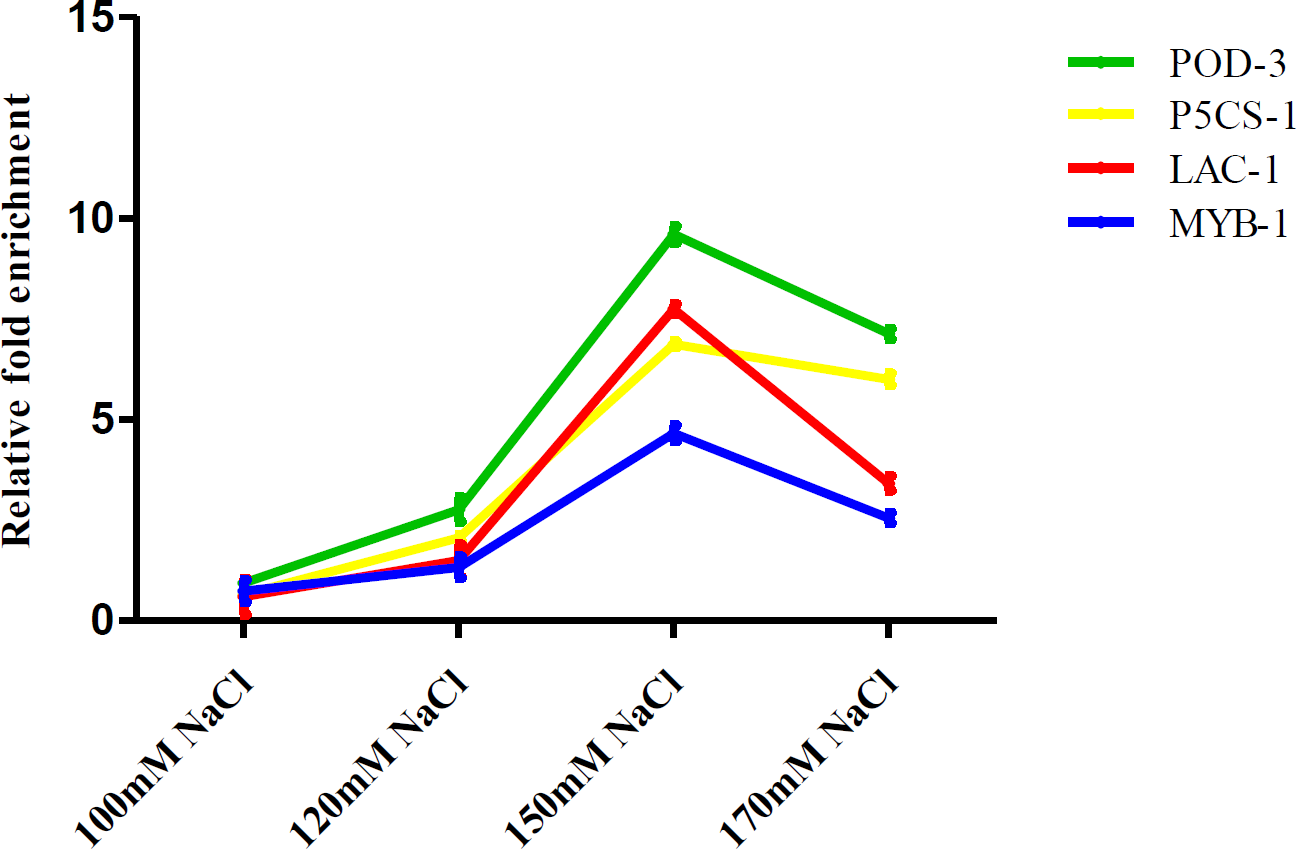
Determination of the optimum NaCl concentration for immunoprecipitation. Different concentrations of NaCl in the OIP (optimized immunoprecipitation) buffer were studied for ChIP immunoprecipitation, and ChIP-qPCR was performed to determine the fold enrichment Four truncated promoters of genes in *B. platyphylla* were studied using ChIP-qPCR. Three independent experiments were performed, and data are means ± SD from three replicates.

### Sucrose plays an important role in improvement of immunoprecipitation

To determine whether sucrose is involved in immunoprecipitation efficiency, OIP buffers containing 0, 5, 7, 9, and 11% (w/v) sucrose were made, and immunoprecipitation was performed. ChIP-qPCR analysis was used to determine the immunoprecipitation efficiency. The results showed that sucrose at 5–11% could improve the efficiency of immunoprecipitation (Fig. 5). In addition, 7% sucrose had the largest effect on immunoprecipitation efficiency, producing an enrichment of 3.54–7.41-fold compared with that of the control (Fig. 5). Therefore, sucrose plays an important role in immunoprecipitation improvement, and 7% sucrose was identified as the most suitable sucrose concentration for immunoprecipitation.

**Figure 5.**
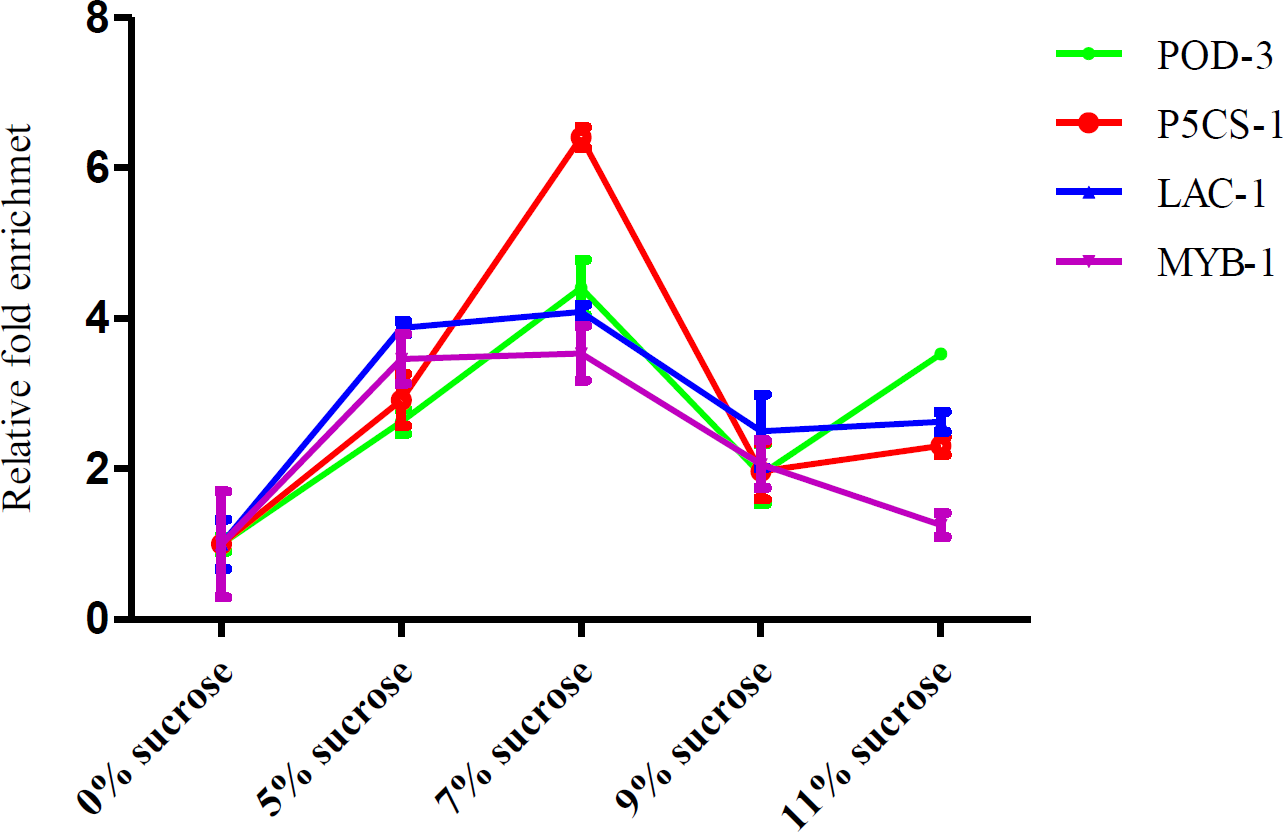
Determination of the effects of sucrose on immunoprecipitation efficiency. Different concentrations of sucrose were added into the OIP (optimized immunoprecipitation) buffer, and immunoprecipitation was performed. Con: Fold enrichment of ChIP performed using OIP buffer without sucrose (control). 5, 7, 9 and 11% sucrose: fold enrichment of ChIP performed using 5, 7, 9 and 11% of sucrose in the OIP buffer. The relative fold enrichment was calculated as 5, 7, 9 and 11%/Con. Four truncated promoters of genes directly regulated by *BplMYB46* were studied in *B. platyphylla* using ChIP-qPCR. Three independent experiments were performed, and data are means ± SD from three replicates.

### Optimization of the crosslinking reversal step

Next, we optimized the procedure for reversing the crosslinking between chromatin and proteins. The ChIP procedure was performed according to the standard ChIP protocol. After elution of the ChIP DNA, the elution product was divided into two equal portions. Proteinase K was added to one portion for crosslinking reversal, and the other portion was reverse crosslinked using NaCl overnight (standard protocol). The ChIP-qPCR results showed that the enrichment of ChIP DNA using Proteinase K digestion increased by 2.41–3.20-fold compared with that achieved using the NaCl-mediated classic crosslinking reversal procedure (Fig. 6).

**Figure 6.**
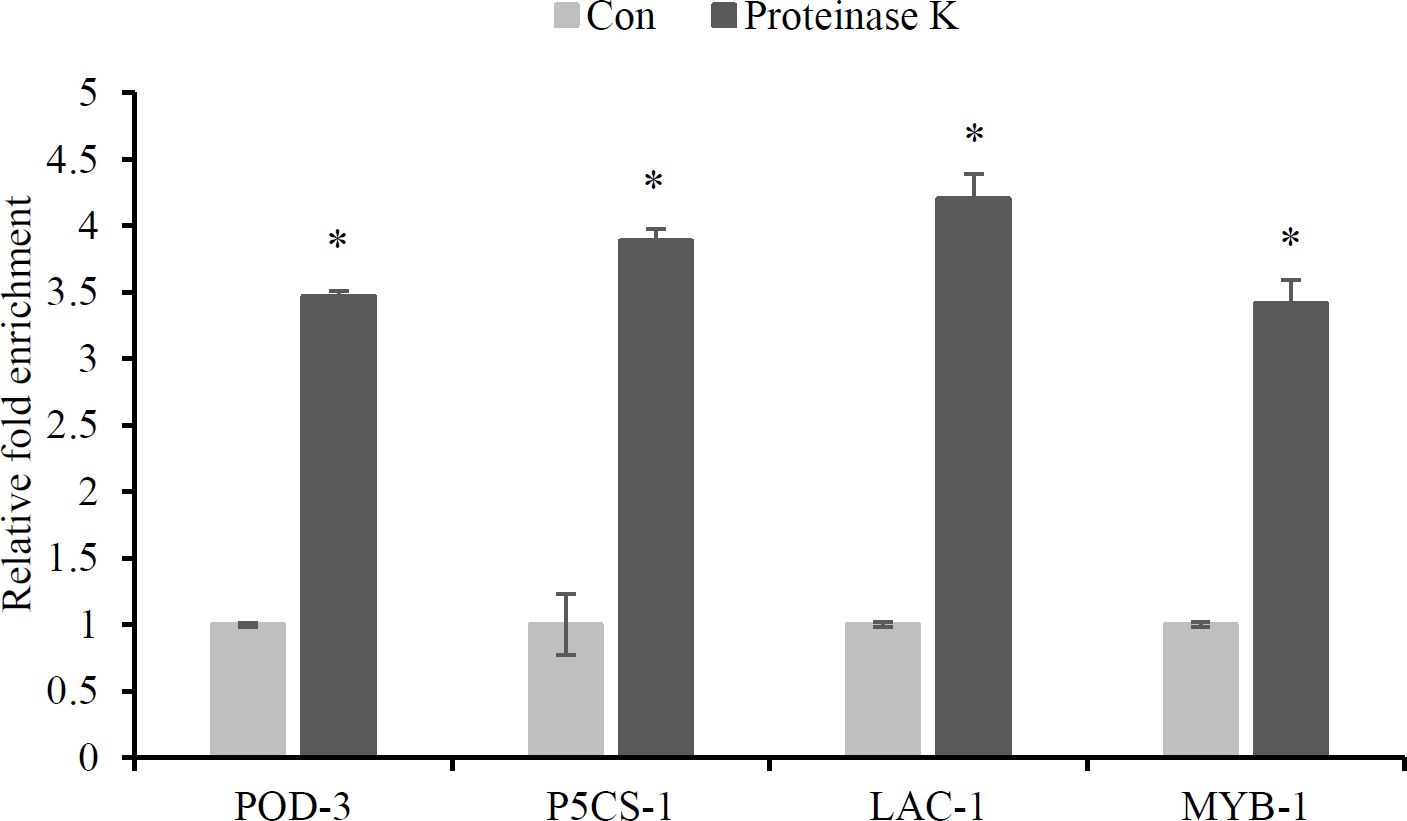
Comparison of methods of crosslinking reversal in ChIP. ChIP was performed according to the classic protocol and the eluted ChIP DNA was divided equally into two portions, which were used for crosslinking reversal using two methods. Method 1: Proteinase K direct digestion to substitute for crosslinking reversal; method 2: crosslinking reversal using NaCl at 65 °C overnight. ChIP-qPCR was conducted to determine the amounts of ChIP DNA. Proteinase K: Fold enrichment of ChIP performed using method 1; Con: Fold enrichment of ChIP performed using method 2 as the control. The relative ChIP fold enrichment was calculated as: Proteinase K/Con. Four truncated promoters of genes directly regulated by *BplMYB46* were studied in *B. platyphylla* using ChIP-qPCR. Three independent experiments were performed, and data are means ± SD from three replicates.

We further compared three different crosslinking reversal methods: (1) proteinase K direct treatment; (2) reverse crosslinking using NaCl; and (3) proteinase K treatment after reversing crosslink using NaCl. Tris-phenol and chloroform extraction was performed to remove the crosslinked DNA, and agarose gel electrophoresis was used to monitor the amount of DNA released by crosslinking reversal. The results showed that crosslinking reversal using NaCl overnight could not completely de-crosslink the DNA; however, direct proteinase K digestion and proteinase K digestion after NaCl crosslinking reversal could completely de-crosslink the DNA (Fig. 7). In addition, compared with the other two approaches, direct proteinase K digestion was a time saving and simpler method (Fig. 7).

**Figure 7.**
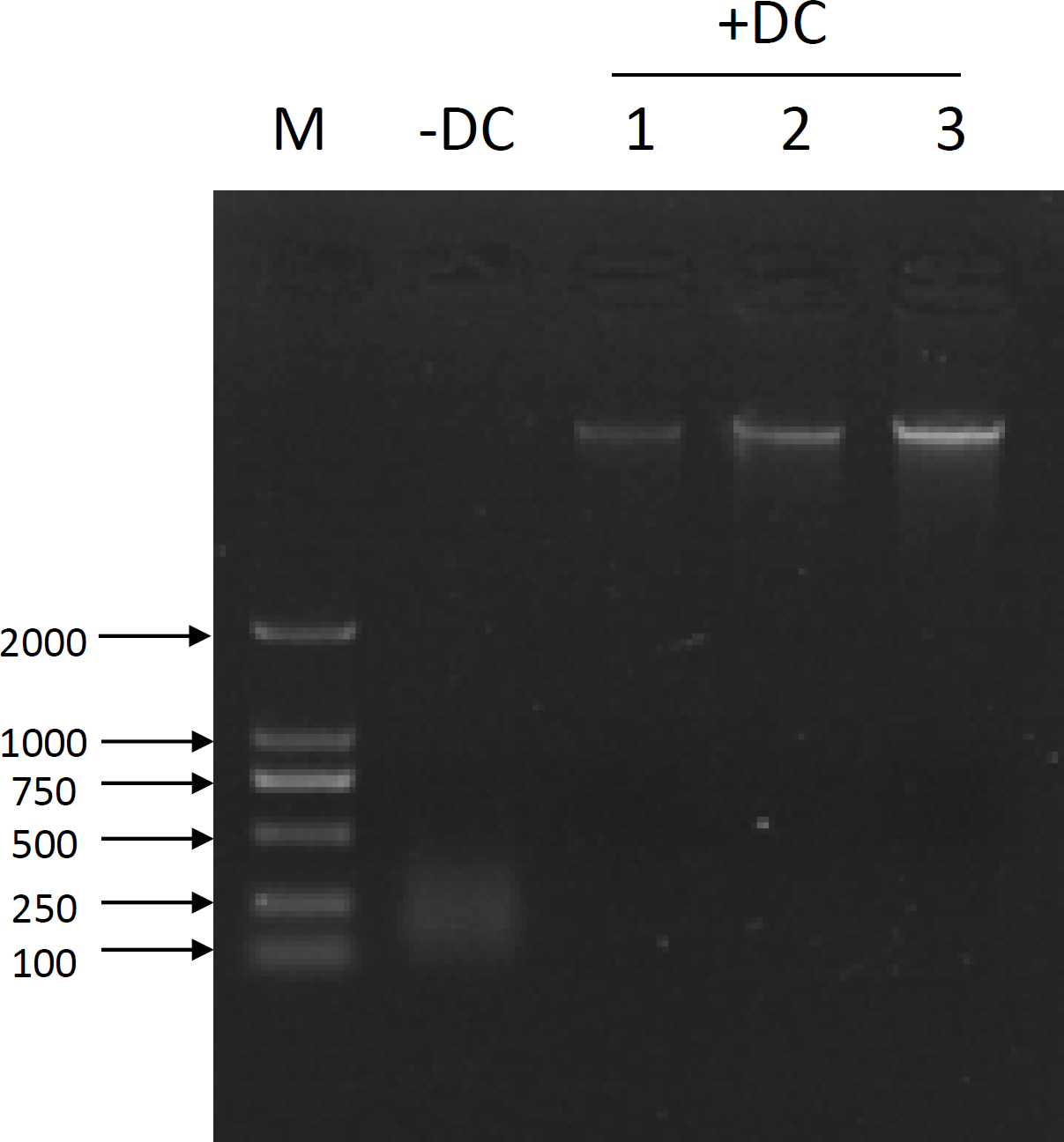
Determination of the efficiency of different crosslinking reversal methods. Three methods were performed on birch chromatin. Method 1: Reversal of crosslinking using NaCl at 65 °C overnight; method 2: Proteinase K digestion after crosslinking reversal using NaCl at 65 °C for 6 h; method 3: Proteinase K direct digestion at 55 °C for 2 h. +DC: samples were decrosslinked. -DC: samples were not decrosslinked. After these three methods were performed, the chromatin was extracted using one volume of Tris-phenol and chloroform (1:1 v/v), followed by extraction using one volume of chloroform. The supernatant was electrophoresed through an agarose gel to determine the quantity of decrosslinked DNA. M: DNA marker; Line 1, 2, 3: Crosslinking reversal of chromatin using methods 1, 2, and 3, respectively.

### Building an improved ChIP protocol and determination of its immunoprecipitation efficiency

Based on the above results, we developed a new ChIP protocol. The procedures are shown in Figure 8. The first improvement is that 3% formaldehyde was used for chromatin crosslinking instead of 1% formaldehyde, which would reduce the decrosslinking caused by sonication. The second improvement is that a protein centrifugal filter was used to concentrate the chromatin. The third improvement was the use of OIP buffer to substitute for classic ChIP Ab incubation buffer for immunoprecipitation. The fourth improvement was the addition of sucrose to increase immunoprecipitation. Finally, in the crosslinking reversal step, proteinase K was used to directly digest proteins, which takes no more than 2 h and achieves complete reversal of DNA crosslinking (Fig. 8).

**Figure 8.**
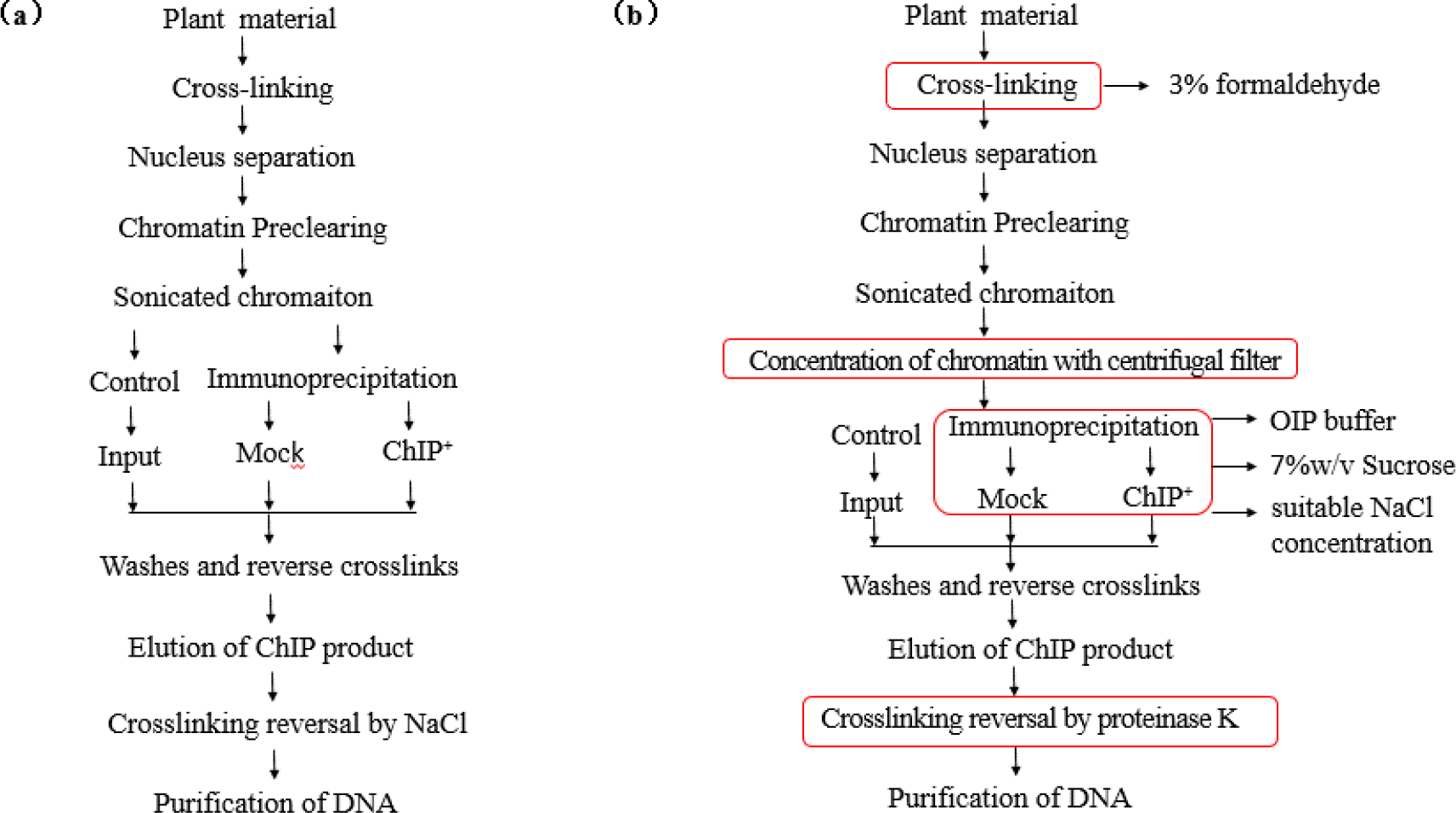
The procedure for the newly developed ChIP protocol. The outline of the developed ChIP. a: The procedures for standard ChIP; b: The procedures for the newly developed ChIP. The optimized procedures are marked with red frames, which included crosslinking protein and chromatin using 3% formaldehyde instead of 1% formaldehyde; purification and concentration of chromatin using protein centrifugal filters; using OIP (optimized immunoprecipitation) buffer instead of standard ChIP Ab incubation buffer for immunoprecipitation; sucrose was added to improve the immunoprecipitation efficiency, and crosslinking reversal was achieved using proteinase K directly. The detailed procedures are shown as supplementary file 1.

Following this improved ChIP protocol, we determined its ChIP efficiency. ChIP-qPCR showed that the fold enrichment using the improved ChIP method was 5.43–20.53-fold (average = 16.88-fold) higher than that achieved using standard ChIP (Fig. 9). Consistently, repeating the experiment in Arabidopsis plants also showed an enrichment of 4.31–9.39-fold (average = 6.43-fold) using the improved ChIP method compared with that achieved using the standard ChIP method (Fig. 9a, b). In addition, the negative control genes were not enriched using the improved ChIP protocol compared with that gained using the standard ChIP method, suggesting that the improved ChIP method results in a lower background (Fig. 9a, b). At the same time, when using the transgenic Arabidopsis plants expressing GFP gene only as material, no fold enrichment of the aim DNA was observed (Fig. 9c), suggesting that this ChIP method can specially enrich aim DNA. Taken together, these results indicated that the improved ChIP procedure improved the efficiency of ChIP markedly.

**Figure 9.**
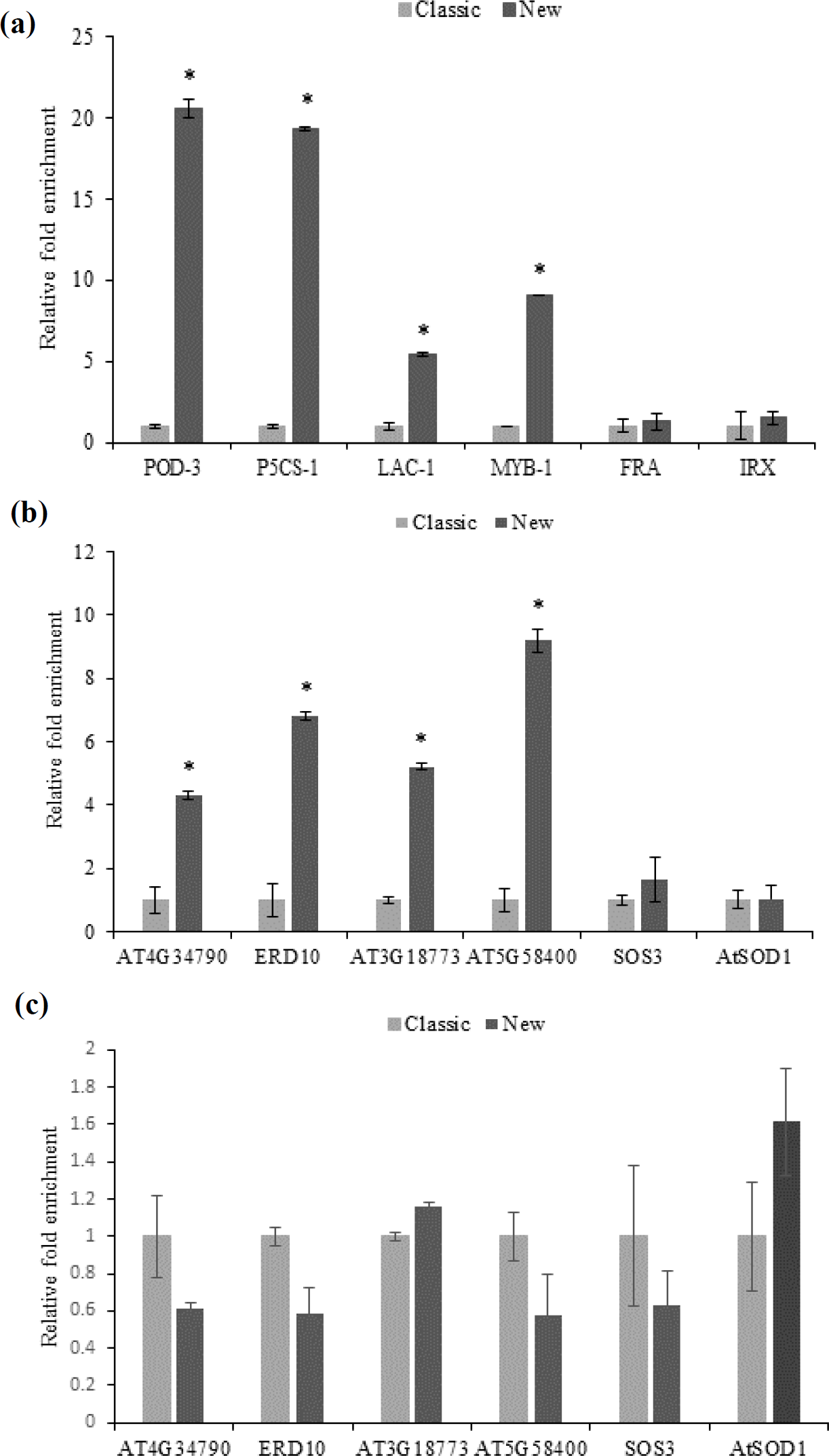
Analysis of the developed ChIP protocol for ChIP enrichment. ChIP was carried out using the improved ChIP protocol and the standard ChIP protocol. The fold enrichment of ChIP was studied using ChIP-qPCR. Classic: fold enrichment of ChIP performed using the standard ChIP protocol; New: fold enrichment of ChIP performed using the new ChIP protocol. The relatively ChIP fold enrichment were calculated as: New/Classic. (a, b) The ChIP fold enrichment values were compared between the classic and the new ChIP protocol in *B. platyphylla* (a) and *A. thaliana* (b). c: The Arabidopsis plants overexpression of GFP were used as material for ChIP, and immunoprecipitation was performed with antiGFP antibidy (as negative control). Four truncated promoters of genes from *B. platyphylla* and four truncated promoters of genes from *A. thaliana* were analyzed using ChIP-qPCR. Three independent experiments were performed, and data are means ± SD from three replicates.

## DISCUSSION

We studied the factors that influence immunoprecipitation efficiency in ChIP, and then built a robust ChIP protocol, which could significantly improve the enrichment of ChIP DNA compared with that achieved using the standard ChIP protocol (Fig. 9).

Compared with the commonly used standard ChIP protocol and the protocol of Li et al. (2014), this procedure has the following differences. (1) The use of 3% formaldehyde for crosslinking to reduce de-crosslinking during sonication; (2) concentration of crosslinked-chromatin using a centrifugal filter before immunoprecipitation; (3) adding sucrose to increase the efficiency of the interaction between the antibody and the antigen; (4) using a more suitable buffer for immunoprecipitation; and (5) recovery of DNA from crosslinked chromatin using proteinase K digestion instead of NaCl reversal.

### Concentration of crosslinked-chromatin is important for immunoprecipitation

In the present study, protein ultrafiltration was employed to concentrate the crosslinked-chromatin, which has the following four advantages: (1) This procedure could eliminate some components released from cells that might inhibit the interaction between the antigen and the antibody, and could also completely remove SDS that will inhibit immunoprecipitation; (2) protein ultrafiltration can change the lysis buffer for the buffer that is most suitable for the antigen–antibody interaction, which would greatly increase the efficiency of immunoprecipitation (Fig. 2); (3) the small molecular weight (<100 kDa) proteins that do not cross-link with chromatin are eliminated using ultrafiltration, which could reduce the background of immunoprecipitation; and (4) this process can adjust the concentration of crosslinked-chromatin to a suitable level for immunoprecipitation. By contrast, it is difficult to solve the above four problems in the standard ChIP protocol. The results showed that concentration of crosslinked chromatin using protein centrifugal filters could significantly increase the efficiency of immunoprecipitation (Fig. 2). In addition, in the present study, the concentration of crosslinked chromatin might not have been the most appropriate for immunoprecipitation, and further adjustments to this concentration might increase immunoprecipitation efficiency.

### Proteinase K digestion is efficient in rescue DNA from crosslinked chromatin

Incubation of crosslinked chromatin in 0.2 M NaCl solution at 65 °C overnight is a commonly used method to reverse DNA crosslinking. Some ChIP procedures also use proteinase K for digestion after incubation in 0.2 M NaCl solution at 65 °C overnight or for at least 6 h (Li *et al.*, 2014, Tsugama *et al.*, 2013, Haring *et al.*, 2007). In the modified procedure, we used proteinase K to directly digest proteins and recover DNA, which could be completed within 2 h. ChIP-qPCR showed that this method increased the enrichment of ChIP products compared with using NaCl to reverse crosslinking (Fig. 6), and has similar efficiency in DNA recovery compared with proteinase K treatment after NaCl incubation (Fig. 6). However, direct proteinase K treatment could save processing time. This method is simple and time saving; therefore, it could be used commonly to reverse crosslinking in many experiments.

### OIP buffer can significantly improve immunoprecipitation efficiency

The interactions between the antigen and the antibody is the key procedure in ChIP; therefore, its optimization is important. In many ChIP protocols, the pH value of the immunoprecipitation buffer is 8.0, which is higher than the pH of the nucleus; therefore, it might not be most appropriate for the immune interaction. In this study, we employed HEPES-NaOH in the OIP buffer to achieve a pH value of 7.5, which is close to the pH value in the nucleus according to Shen et al (2013). Moreover, some reagents in the buffer were also optimized. Experiments showed that the OIP buffer could improve the immunoprecipitation efficiency greatly compared with using the ChIP Ab incubation buffer from the in standard ChIP protocol (Fig. 3). Therefore, the OIP buffer is more suitable for use in ChIP in plants than the ChIP Ab incubation buffer.

### Sucrose is important in improvement of immunoprecipitation

To improve the efficiency of immunoprecipitation, we further studied other reagents, including sucrose, trehalose, PEG, and glycerol (data not shown). Among them, only sucrose displayed significantly increased efficiency of immunoprecipitation. In addition, the results showed that 5–11% sucrose could improve immunoprecipitation (Fig. 5). The role of sucrose in improving immunoprecipitation has not been reported previously. Thus, to increase immunoprecipitation efficiency sucrose could be used in experiments involving interactions between the antibody and the antigen.

### The reason for using birch plants as researching material

In the present study, we use birch as the plant material for ChIP investigation. Birch is a woody plant species that is distributed widely in the cold temperate zone from Europe to Asia, and shares many characteristics with other woody plants in terms of its structure and composition. In addition, birch contains substantially higher levels of chemical components compared with other woody plants, including polysaccharides and polyphenols, which make it difficult to isolate DNA, RNA, or protein from this plant (especially from the mature leaves of birch) compared with many other woody plant species. Our developed ChIP protocol works well in birch; therefore, it should also work well in other woody plant species. The improved ChIP works well in birch and Arabidopsis; therefore, it could be adapted for use in other woody and herbaceous plants.

We should note that ChIP efficiency was improved more in birch than in Arabidopsis using the improved ChIP protocol (Fig. 9). This might be explained by the fact that birch has many kinds of metabolites, such as polysaccharides and polyphenols, which will reduce immunoprecipitation efficiency quite highly than Arabidopsis plants, leading to the standard ChIP procedure not working well. Therefore, there is a large scope for improvement using the newly developed ChIP protocol. However, in Arabidopsis, there are few metabolites that hinder the immunoprecipitation efficiency compared with that in birch, and the standard ChIP could work well, resulting limited improvement in ChIP efficiency compared with that achieved in birch. Therefore, this improved ChIP protocol might be more useful in the plants that have abundant metabolites that hinder immunoprecipitation efficiency.

## CONCLUSIONS

In the present study, an improved ChIP protocol was developed that includes five improved procedures: (1) Crosslinking proteins and chromatin using 3% formaldehyde instead of 1% formaldehyde; (2) concentration of crosslinked chromatin; (3) using an optimized IP buffer; (4) the addition of sucrose to improve immunoprecipitation efficiency; and (5) the use of proteinase K to digest proteins crosslinked with DNA (Fig. 8). Improvement of each of the above procedures could increase the enrichment of ChIP DNA significantly. Together, these five improvements could increase the enrichment of ChIP DNA greatly. In addition, this study might also provide helpful guidance to improve other experiments involving immunoprecipitation.

## MATERIALS AND METHODS

### Plant materials and the DNA sequences used in ChIP

Two kinds of plant species were used as research materials, i.e. a woody plant species, birch (*B. platyphylla*), and a herbaceous plant species, *A. thaliana*. Four genes that had been identified to be directly regulated by BplMYB46 in birch (Guo *et al.*, 2017), and birch plants overexpressing BplMYB46-FLAG (*BplMYB46*, Genbank number: KP711284) were used in the ChIP experiments. Four genes that were directly regulated by AST1 (*AST1*, AT3G24860) in *A. thaliana*(Xu *et al.*, 2018), and the Arabidopsis plants overexpressing AST1-GFP were also used.

### Procedure for standard ChIP

The ChIP procedure followed that of Haring et al (2007) with some modifications. The detailed procedures were as follows: ***Plant material and crosslinking*:** (1) One gram of plant sample (aerial part; for birch sample used that is 5cm in height; for Arabidopsis sample, 4-week-old T3 transgenic plants were used) was incubated in 30 ml of buffer A to cross-link the protein and DNA under vacuum conditions for 10 min. Then, the sample was added with 2.5 ml of 2 M glycine, mixed well, and incubated for 5 min to stop the cross-linking reaction. The solution was removed from the sample, which was washed twice with cold milliQ water, and excess moisture was thoroughly removed using paper towels. ***Nuclei isolation*:** (2) The samples were ground to a fine powder under liquid nitrogen, add with 30 ml of buffer B, and mixed well. The following procedures were all performed on ice. (3) The solution was filtered through four layers of Miracloth, and the filtrate was collected into a new 50 ml tube, and centrifuged for 20 min at 2,800 × *g* at 4 °C. The precipitates were resuspended in 20 ml of buffer C, and the solution was centrifuged for 10 min at 12,000 × *g* and 4 °C; this step was repeated until the precipitate became white. (4) The precipitate was resuspended in 300 μl of buffer D as the sample solution. Then, 600 µl of buffer E was added to a new 1.5 ml tube, and overlaid with the sample solution before centrifugation at 12000 × *g* for 45 min at 4 °C. ***Chromatin sonication*:** (5) The supernatant was removed and the pellet was resuspend in 320 µl of lysis buffer; a 10 µl aliquot sampled as “unsheared chromatin”. (6) The chromatin solution was sheared into 200-to 1000-bp fragments using an ultrasonic homogenizer (Scientz-IID, Scientz Biotechnology, Ningbo, China) with the following parameters: Sonication for 3 sec and stopping for 15 sec, at 10% power setting for a total of 20 min. (7) The sonicated chromatin was centrifuged at 12,000 × *g* for 5 min at 4 °C. A 10 µl aliquot of the chromatin solution was sampled to check the sonication efficiency. Then, a 50-µl aliquot of the supernatant was reserved as the Input sample. ***Chromatin preclearing*:** (8) Then, 200 µl of the supernatant was diluted 10-fold by adding 1.8 ml of ChIP Ab incubation buffer. Washed and blocked protein A/G Agarose beads (40 µl) were added to the chromatin solution and the mixture was incubated for 1 h at 4 °C with gentle agitation (12 rpm). The chromatin solution was then centrifuged at 1000 × g at 4 °C for 5 min to pellet the beads. ***Preparation of washed and blocked protein A/G agarose beads*:** (9) The supernatant of the protein A/G agarose beads (150 µl) was removed by centrifugation at 1000 × *g* for 5 min, and 1 ml of ChIP Ab incubation buffer was added. The beads were centrifuged at 1000 × *g* for 5 min to precipitate the beads. This rinsing process was repeated twice. The beads were resuspended in 150 µl of ChIP Ab incubation buffer with bovine serum albumin (BSA) to a final concentration of 10 µg/ml. ***Immunoprecipitation*:** (10) Two portions of 600 µl of the chromatin solution from step 8 were taken, one portion was added with 6 µl of anti-GFP or anti-FLAG antibodies as the ChIP sample (ChIP+), the other was added with 6 µl anti-HA antibody as the mock control (ChIP-), and both were incubated at 4 °C overnight. (10) Washed protein A/G Agarose beads (60 µl) were added to the two tubes, respectively. The tubes were incubated for 3 h at 4 °C with gentle agitation. (12) The tubes were centrifuged for 5 minutes at 1000 × g at 4 °C to precipitate the beads. ***Washing*:** (13) The beads were washed for 10 min sequentially with 1 ml of the following buffers: Low Salt Buffer, high salt buffer, LiCl wash buffer, and TE buffer. (14) Then, 250 µl of prewarmed (65 °C) elution buffer (50mM Tris pH 8.0, 10 mM EDTA, 1% SDS) was added, and the beads were resuspended by tapping and incubated at room temperature for 15 min (mixing at 5-min intervals). The beads were centrifuged for 5 minutes at 2000 × *g*, the supernatant was transferred into a new tube, the elution step was repeated once, and the two eluates were mixed together. ***Reverse crosslinking*:** (15) 5 M NaCl solution was added to the IP or mock control elution to achieve a concentration of 0.2 M NaCl. Then, 100 µl of TE buffer was added to the input sample, and 5 M NaCl was added to a concentration of 0.2 M NaCl. All samples were reverse cross-linked overnight at 65 °C. ***DNA purification*:** (16) The reverse crosslinked samples were purified using a DNA purification spin column (Qiagen, Hilden, Germany), and the column was eluted twice with 60 µl of TE buffer. Fold enrichment calculation was following Haring et al (2007).

### Analysis of chromatin decrosslinking caused by sonication

To investigate the effects of different concentrations of formaldehyde on chromatin decrosslinking caused by sonication, birch plant samples were crosslinked with 1% or 3% formaldehyde and treated by sonication as described in “Procedure for standard ChIP”. After sonication, the decrosslinked chromatin was harvested by extraction with an equal volume of Tris-phenol and chloroform, and the supernatant was further extracted using an equal volume of chloroform. Then, the decrosslinked DNA was harvested from the supernatant using a PCR purification kit (Qiagen). The extent of DNA decrosslinking was analyzed using quantitative PCR.

### Analysis of the effects of concentration of crosslinked chromatin

To concentrate the crosslinked chromatin, the chromatin was sonicated, centrifuged (according to step 7 in the above ChIP protocol), and diluted 10-fold by adding ChIP Ab incubation buffer (which dilutes the SDS to 0.1% to avoid micelle formation, which would affect protein ultrafiltration), and the sonicated chromatin was purified using protein centrifugal filters (30 kDa cutoff, Millipore, Billerica, MA, USA) to 300 µl, then added with 2700 µl of ChIP Ab incubation buffer, mixed well, and concentrated with centrifugal filters to 300 µl again (following its manufacturer instructions). The purified chromatin was divided into two equal portions. One was used for immunoprecipitation (ChIP+) and the other was used as a no antibody immunoprecipitation (ChIP-) control. The same volume of antibody as was used in the classic protocol was added for immunoprecipitation. The subsequent steps were same as those in the standard ChIP procedure. ChIP-qPCR was performed to study the effects of protein centrifugal filter concentration.

### Analysis of the ChIP efficiency using an optimized immunoprecipitation buffer

The OIP buffer (20 mM HEEPS-NaOH pH 7.5, 150 mM NaCl, 1 mM EDTA, 1% TritonX-100, 1 mM PMSF, proteinase inhibitors at 1 µg/ml each) was used to replace the ChIP Ab incubation buffer in the immunoprecipitation step. Other procedures and buffers were the same as those in the classic procedure described in above. ChIP-qPCR was used to compare the immunoprecipitation efficiency between using OIP and ChIP Ab incubation buffers.

### Analysis of the effects of different concentrations of NaCl in OIP buffer

To determine the optimal concentration of NaCl for immunoprecipitation, NaCl in the OIP buffer was used at 110, 120, 150, and 170 mM, and other reagents in OIP buffer were unchanged. Other procedures and buffers were the same as the classic procedure described in above. ChIP-qPCR was performed to investigate the most suitable NaCl concentration for immunoprecipitation.

### Investigation of the effect of sucrose on immunoprecipitation

To determine the effect of sucrose on immunoprecipitation, sucrose at 5, 7, 9 and 11% (w/v) were added into the OIP buffer for immunoprecipitation. The other steps of ChIP followed the standard ChIP protocol. ChIP-qPCR was performed to study whether sucrose at different concentrations affected ChIP immunoprecipitation.

### Comparison of different methods of crosslink reversal

Two methods to recover DNA from crosslinked chromatin were compared. After the ChIP product elution (step 13), the eluted product was divided into two equal portions. Portion one was added with Proteinase K to a final concentration of 0.7 μg/μl and incubated at 55 °C for 2 h to decompose the crosslink between the protein and chromatin. Portion two was subjected to chromatin crosslinking reversal following the classic procedure, i.e., added with NaCl to a final concentration of 0.2 M, and incubated at 65 °C overnight.

Then, DNA was purified from these two portions using a DNA purification spin column (Qiagen), and eluted with 60 µl of TE buffer. To compare these two methods of recovery of DNA from crosslinked chromatin, qPCR was performed.

### Comparison of the methods to recover DNA from crosslinked chromatin

Three methods to purify DNA from crosslinked chromatin were compared. After elution of the ChIP products (step 13), the eluted products were divided into three equal portions. Method one used reverse crosslinking from the classic method, i.e. following the procedure described at step 14 (as a control). In method two, the sample was added with proteinase K to a final concentration of 0.7 μg/μl and incubated at 55 °C for 2 h. In method three, crosslinking reversal was performed according to the method of Li et al (2014), i.e. NaCl to the final concentration of 0.2 M was added, and the sample was incubated at 65 °C for 6 h to reverse cross-linking. Then, 32.5 μl of protease/RNase buffer (150 mM EDTA (pH 8.0), 615 mM Tris-HCl (pH 6.5), 14 mg/ml proteinase K, and 0.30 μg/μl RNase) was added and the mixture was incubated at 45 °C for 1 h.

The products from these three methods were extracted separately with an equal volume of Tris-phenol (pH 8.0) and chloroform (1:1 v/v), and then extracted with an equal volume of chloroform. Ten microliters of supernatant was electrophoresed through an agarose gel to determine the quantity of DNA.

### The protocol of the newly developed ChIP technique

The improved ChIP protocol was developed (Fig. 6) using the following procedures. Step 1 used 3% formaldehyde instead of 1% formaldehyde for chromatin crosslinking. Step 2–7 were the same as classic protocol described above. In step 8, 200 µl of the supernatant was diluted 10-fold by adding 1.8 ml of OIP buffer (supplemented with 7% sucrose), and was concentrated using protein centrifugal filters (30 kDa cutoff) to 300 µl. Then, a 2700 µl of OIP buffer (supplemented with 7% sucrose) was added and mixed well. The sample was then concentrated using centrifugal filters to 400 µl. In step 10, two portions of 200 µl of the chromatin solution from step 8 were taken. One portion was added with 6 µl of the target antibody as the ChIP sample (ChIP+), the other portion had antibody added (or was added with another antibody) as the mock control (ChIP-), and incubated at 4 °C overnight. Steps 11–14 followed the classic protocol. In step 15, the elution buffer was added with proteinase K to a final concentration of 0.2 μg/μl and incubated at 55 °C for 2 h. Step 16 was the same as the classic protocol. The detailed protocol of developed ChIP was shown as Supplementary file 1.

### Quantitative real-time PCR analysis

Quantitative real-time PCR was performed on a qTower 2.2 (Analytik Jena AG, Jena, Germany). The PCR reaction system contained 10 µl of SYBR Green Real-time PCR Master Mix (Toyobo, Osaka, Japan), 0.5 µM of each forward and reverse primes, and 2 µL of ChIP product as the PCR template, with a total volume of 20 µl. The PCR reaction was conducted with the following parameters: 94 °C for 1 min; 40 cycles of 94 °C for 12 s, 58 °C for 30 s, and 72 °C for 45 s. Melting curves were generated for each reaction to identify the specificity of the PCR reaction. The DNA sequence of ubiquitin (FG065618) and ACT7 (AT5G09810) were used as the internal controls in birch and Arabidopsis, respectively. The primers for ChIP-PCR used in birch were the same as those used by Guo et al (2017), and the primers used in Arabidopsis were the same as those used by Xu et al (2018). The primers used for ChIP-qPCR are shown in Table S1.

### Reagents

Sucrose (Sigma-Aldrich); Formaldehyde solution 37% (wt/wt) (Sigma-Aldrich); Glycine (Fisher Scientific); Tris base (Promega); Hydrochloric acid (HCl; Sigma-Aldrich); PMSF, aprotinin, leupeptin, and pepstatin A (Sigma-Aldrich); DMSO (Sigma-Aldrich); β-Mercaptoethanol (Sigma-Aldrich); Miracloth (Calbiochem, San Diego, CA, USA); anti-FLAG antibody (Sigma-Aldrich, SAB4301135); anti-GFP antibody(Sigma-Aldrich, SAB4301138); protein A/G Agarose beads (Beyotime Biotechnology, Shanghai, China); Triton X-100 (Solarbio); EDTA disodium salt (Solarbio); Proteinase K (Promega); RNase A solution (Promega).

### ChIP Buffers

Buffer A comprised 10 mM Tris-HCl (pH 8.0), 0.4 M sucrose, 1.0 % w/v formaldehyde, 1 mM PMSF, proteinase inhibitors (aprotinin, leupeptin, pepstatin A, 1 µg/ml each), and 5 mM β-mercaptoethanol.

Buffer B comprised 10 mM Tris-HCl (pH 8.0), 0.4 M sucrose, 5 mM MgCl_2_, 5 mM β-mercaptoethanol, 1 mM PMSF, and proteinase inhibitors (aprotinin, leupeptin, pepstatin A) at 1 µg/ml each.

Buffer C comprised 10 mM Tris-HCl (pH 8.0), 0.25 M sucrose, 10 mM MgCl_2_, 1.2% Triton X-100, 5 mM 2-mercaptoethanol, 1 mM PMSF, and proteinase inhibitors (aprotinin, leupeptin, pepstatin A) at 1 µg/ml each.

Buffer D comprised 10 mM Tris/HCl pH 8.0, 0.7 M sucrose, 2 mM MgCl_2_, 5 mM 2-mercaptoethanol, 1 mM PMSF, and proteinase inhibitors (aprotinin, leupeptin, pepstatin A) at 1 µg/ml each.

Buffer E comprised (10 mM Tris/HCl pH 8.0, 2.0 M sucrose, 2 mM MgCl_2_, 5 mM 2-mercaptoethanol, 1 mM PMSF, and proteinase inhibitors (aprotinin, leupeptin, pepstatin A) at 1 µg/ml each.

Lysis buffer comprised 50 mM Tris pH 8.0, 10 mM EDTA, 1 % w/v SDS, 1 mM PMSF, and proteinase inhibitors (aprotinin, leupeptin, pepstatin A) at 1 µg/ml each.

ChIP Ab incubation buffer comprised 20 mM Tris pH 8.0, 1.2 mM EDTA, 1.2% v/v Triton X-100, 150 mM NaCl, 1 mM PMSF, and proteinase inhibitors (aprotinin, leupeptin, pepstatin A) 1 µg/ml each.

Low Salt Buffer comprised 20 mM Tris pH 8.0, 2 mM EDTA, 0.1% w/v SDS, 1% v/v Triton X-100, and 150 mM NaCl).

High salt buffer comprised 20 mM Tris-HCl, pH 8.0, 500 mM NaCl, 0.1% SDS, 1% Triton X-100, and 1 mM EDTA) (wash three times);

LiCl wash buffer comprised 25 mM LiCl, 0.1% sodium deoxycholate, 1 mM EDTA, and 20 mM Tris-HCl, pH 8.0 (two washes).

TE buffer comprised 10 mM Tris-HCl, pH 8.0, and 1 mM EDTA (two washes). Elution buffer comprised 50 mM Tris pH 8.0, 10 mM EDTA, and 1% SDS.

## ACKNOWLEDGMENTS

Not applicable.

## CONSENT FOR PUBLICATION

Not applicable.

## Supplemental materials

**Supplemental file 1**. The procedure of new developed ChIP.

**Supplemental Table 1.** The primers used in ChIP-qPCR

